# Parallel in-depth analysis of repeat expansions: an updated Clin-CATS workflow for nanopore R10 flow cells

**DOI:** 10.1101/2024.11.05.622099

**Authors:** Veronika Scholz, Veronika Schönrock, Hannes Erdmann, Vitus Prokosch, Maria Schoedel, Manuela Almus, Mayra Sauer, Veronika Mayer, Eva Breithausen, Inga van Buren, Christine Dineiger, Caroline Heintz, Ariane Hallermayr, Teresa Neuhann, Elke Holinski-Feder, Angela Abicht, Anna Benet-Pagès, Morghan C Lucas

## Abstract

Hereditary ataxias, caused by expansions of short tandem repeats, are difficult to diagnose using traditional PCR and Southern blot methods, which struggle to detect complex repeat expansions and cannot assess repeat interruptions or methylation. We present an updated Clinical Nanopore Cas9-Targeted Sequencing (Clin-CATS) workflow for analyzing repeat expansions, now compatible with the Oxford Nanopore Technologies R10 flow cell. The workflow incorporates the ONT wf-human-variation Epi2Me workflow, including the Straglr tool to analyze basecalled reads, ensuring compatibility with past, current, and future sequencing chemistries. It expands the number of genes analyzed from 10 to 27 within the ataxia panel and introduces new gene panels for myopathy, neurodegeneration, and ALS/motor neuron disease. Validated with Coriell reference and clinical samples, this method improves the analysis of pathogenic repeat expansions, providing deeper insights into repeat structures while addressing the limitations of traditional approaches.

## INTRODUCTION

Hereditary ataxias are a heterogeneous group of neurodegenerative disorders primarily characterized by progressive motor coordination loss, gait disturbances, and various neurological symptoms such as ocular motor dysfunction, dysarthria, and sensory neuropathy^1,2^. A considerable proportion of these disorders are linked to the expansion of short tandem repeats (STRs)—specific sequences of DNA repeated in both coding and non-coding regions of the genome. These repeat expansions interfere with normal gene function and are responsible for several neurodegenerative diseases with varying levels of severity^3,4^. To date, approximately 60 Mendelian disorders are known to be caused by such expansions, including prominent examples like most spinocerebellar ataxias (SCA), Friedreich ataxia (FRDA), and fragile X-associated tremor/ataxia syndrome (FXTAS)^3,4^.

Traditionally, diagnosing these repeat expansion disorders has relied on PCR-based methods and Southern blotting. While these methods have been the gold standard, they face notable limitations, especially in detecting large, complex repeat expansions. They are also unable to capture information on alternative repeat motifs, repeat interruptions, or epigenetic modifications such as DNA methylation, which are critical for accurate diagnosis and prognosis^2,5^. Furthermore, these methods require iterative testing of specific loci, making them both time-consuming and labor-intensive.

With the development of long-read sequencing technologies, particularly those provided by Oxford Nanopore Technologies (ONT), there has been a significant leap forward in the parallel and comprehensive analysis of complex repeat expansions^4,6,7^. Our previous work introduced Clinical Nanopore Cas9-Targeted Sequencing (Clin-CATS), a method that combined ONT’s R9 chemistry with CRISPR-Cas9-based enrichment to analyze 10 repeat loci associated with the most common hereditary ataxias in the European population at that time^8^. Clin-CATS allowed for the precise quantification of repeat length, sequence composition, and methylation status in key genes associated with adult-onset ataxias, including *ATXN1*, *ATXN2*, *ATXN3*, *ATXN7*, *ATXN8*/*ATXN8OS*, *CACNA1A, FMR1*, *FXN, TBP* and *RFC1*^9–11^.

With the recent introduction of ONT’s R10 flow cell chemistry, offering improved accuracy in resolving homopolymer regions and complex repeat structures, we have adapted and updated the Clin-CATS workflow to leverage the advantages of R10 sequencing. The R10 chemistry offers significant improvements in sequencing accuracy and resolving complex genomic structures, particularly for regions with challenging repeat compositions, such as *RFC1*. These enhancements enable more precise determination of repeat sizes and composition, which is crucial for accurate diagnosis and understanding of the genetic underpinnings of ataxia phenotypes. However, despite these technological advances, ONT’s Cas9-targeted sequencing has been discontinued and has not been re-released for use with the R10 flow cell chemistry. This development required us to modify the original protocol to maintain the high precision and diagnostic utility of Clin-CATS.

In this study, we describe modifications to the Clin-CATS library preparation protocol and bioinformatic pipeline necessary to accommodate the changes introduced by the R10 flow cell chemistry. A key improvement in the bioinformatic workflow is the use of the Straglr tool^12^, which was integrated into the ONT wf-human-variation pipeline. Unlike previous tools that relied on the raw electrical signal from the sequencer and needed to be calibrated to the specific flow cell, Straglr processes data from basecalled reads. This approach is crucial, as the raw signal will likely change with future flow cell chemistry updates. By performing the analysis on basecalled data, our bioinformatic pipeline can be consistently applied across old, current, and future flow cell chemistries, ensuring long-term adaptability—a requirement for accredited diagnostic workflows.

In addition to these workflow improvements, we have expanded the number of genes analyzed with Clin-CATS from 10 to 27. This update supplements the ataxia panel and introduces three additional gene panels, each tailored to a specific group of related disorders: a Myopathy panel, a Neurodegeneration panel, and an ALS/motor neuron disease panel. These panels enable the simultaneous investigation of multiple repeat expansion-related genes across various neurological conditions, further increasing the diagnostic utility of the Clin-CATS workflow.

We demonstrate the capability of the updated Clin-CATS workflow in analyzing pathogenic repeat expansions in a cohort of ataxia patients, providing deeper insights into repeat structures and their clinical relevance. Moreover, we utilized Coriell reference samples alongside clinical samples, ensuring the robustness and accuracy of our approach. This updated method continues to offer a comprehensive and scalable approach for diagnosing and understanding adult-onset ataxias while addressing the limitations of traditional PCR-based and Southern blot methods.

## RESULTS

### Expansion of Clin-CATS panels increases diagnostic utility for ataxia, myopathy, neurodegeneration, and ALS

We expanded the Clin-CATS workflow by increasing the number of target genes from 10 to 27 and introducing four distinct gene panels for ataxia, myopathy, neurodegeneration, and Amyotrophic lateral sclerosis (ALS)/motor neuron disease (**Figure 1**, **Supplementary Table 1**). These tailored panels enable the simultaneous investigation of multiple repeat expansion-related genes across various neurological conditions, significantly enhancing the diagnostic utility of Clin-CATS. Several new target regions were identified from the literature, and we designed guide RNAs specifically for these regions (**Supplementary Table 1**). We confirmed that the guide RNAs accurately excise the newly included targets, such as key genes associated with spinocerebellar ataxias (e.g., SCA27B, SCA10, SCA31, and SCA37), motor neuropathies, myopathies, and neurodegenerative disorders like ALS and Huntington disease-like 2, by sequencing and verifying the alignments with the Integrative Genomics Viewer (IGV)^13,14^ (**Figures S1-3**).

**Figure 1.**
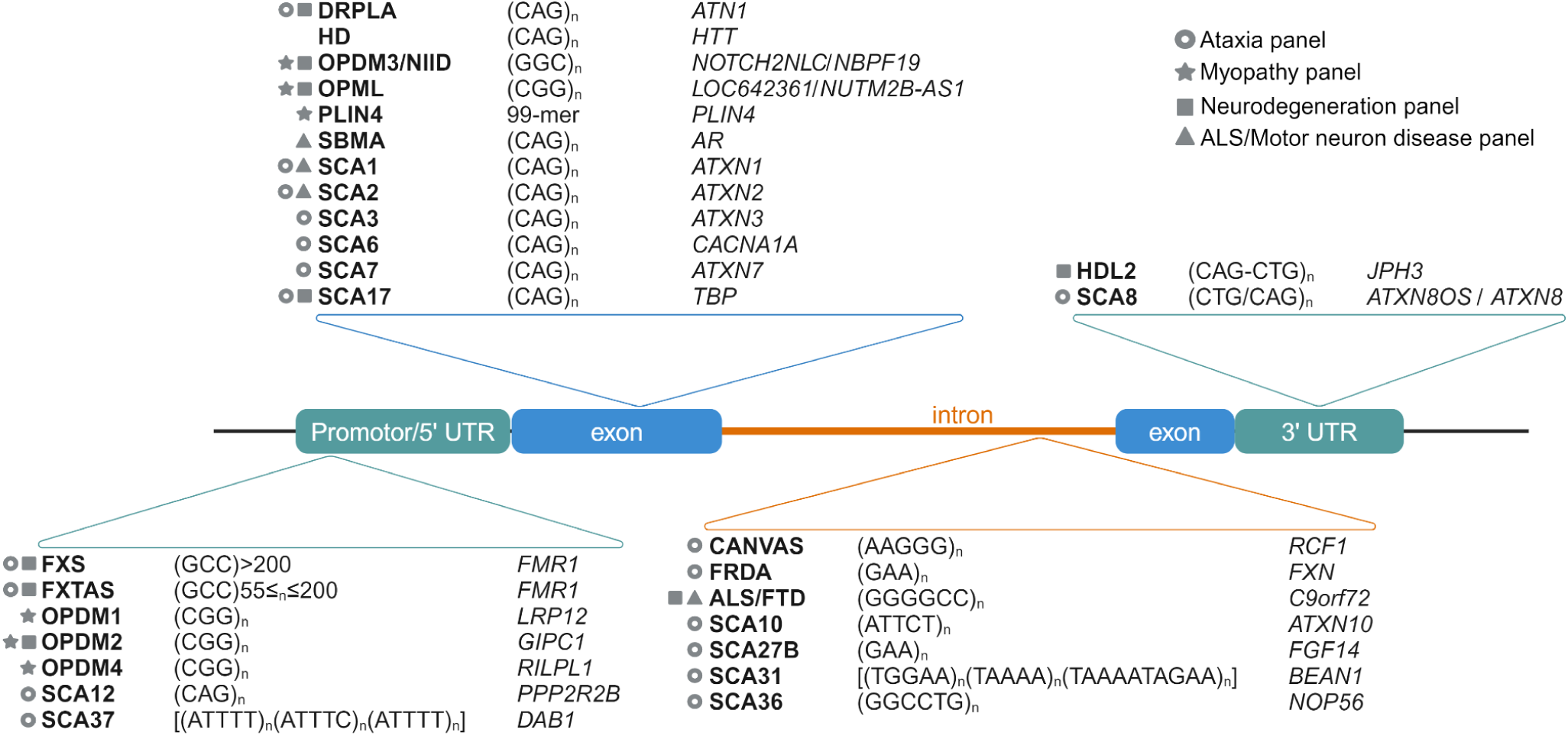
Overview of the updated Clin-CATS target genes and panels. Summary of the disease, repeat motifs, target genes included in the updated Clin-CATS workflow, and their approximate genomic loci (promoter, 5’UTR, exon, intron, 3’UTR). The figure also highlights the gene panels in which each target gene is relevant (ataxia, myopathy, neurodegeneration, and ALS/motor neuron disease). For additional details and complete disorder/condition names, refer to **Supplementary Table 1**.

### R9 protocol on R10 flow cells yields suboptimal coverage for diagnostics

Although ONT has discontinued their Cas9 protocol^15^, similar reagents used in the ONT kit can be purchased directly from Integrated DNA Technologies (IDT) and New England Biolabs (NEB), allowing the continuation of the workflow. The performance of the ONT Cas9 protocol on R10 flow cells resulted in suboptimal coverage, raising concerns about its use in our clinically validated Clin-CATS diagnostic assay. Given that this assay is accredited for clinical diagnostics in Germany, achieving the required quality control metrics on R10 MinION flow cells is essential. In two replicates, only 21/26 and 18/26 target genes reached the required 50X on-target coverage for diagnostics, with 5 and 8 target genes falling short of this threshold (Control condition, **Supplementary Table 2**, **Figure S4**). The percentage of sequenced reads aligning to target regions was 6.9% and 1.3%, respectively (**Supplementary Table 2**). The inability to reach the target coverage on both replicates suggests that the Cas9 protocol for R9 flow cells is not directly transferable to R10 flow cells, as it fails to provide the necessary coverage consistency required for diagnostic accuracy.

### Adaptations to Clin-CATS Cas9 protocol for R10 ensure coverage thresholds for diagnostics

To address the suboptimal performance of the Cas9 protocol on R10 MinION flow cells, we tested three modifications to the default Cas9 library preparation protocol (based on SQK-CS9109): extending the dA-tailing incubation step at 37°C from 15 to 60 minutes (bind60), extending the adapter ligation incubation time from 10 to 30 minutes at room temperature (lig30), and extending the adapter ligation incubation time from 10 minutes at room temperature to overnight at 16°C (ligON). Based on these results, we further evaluated the combination of the first two modifications, testing the impact of simultaneously extending both the dA-tailing and ligation steps (bind60lig30).

Among the tested conditions, the ligON protocol produced the highest number of sequenced reads, with a 9.3-fold increase compared to the control (default Cas9 protocol on an R10 flow cell) (**Supplementary Table 2**). Despite the higher read count, the mean coverage across target regions remained similar to the control, and 5 of the 26 target genes did not achieve the required 50X coverage (**Supplementary Table 2**, **Figure S4**). Furthermore, the percentage of reads aligning to the target regions was 0.3%, indicating a substantial proportion of off-target reads, potentially due to the extended overnight ligation time. Prolonged ligation periods can increase the likelihood of nonspecific ligation events, such as adapter dimer formation or ligation to unintended off-target fragments, reducing the overall efficiency and specificity of the library preparation.

The bind60 and lig30 conditions produced more moderate increases in sequencing reads, with fold changes of 1.6 and 2.0, respectively (**Supplementary Table 2**). Both conditions had improved mean coverage relative to the control, with bind60 achieving a 2.2-fold increase in coverage and lig30 showing a 1.9-fold increase. Importantly, in the bind60 condition, all target genes were covered in both replicates, while in the lig30 condition, one replicate achieved full coverage, and the other missed 4 genes, covering 22 out of 26 (**Supplementary Table 2**, **Figure S4**).

Given the improvements in sequencing metrics observed in bind60 and lig30, we hypothesized that combining these adaptations might yield further enhancements. In the bind60lig30 condition, the average number of reads was 1.7-fold higher than the control, while mean coverage increased by 2.0-fold (**Supplementary Table 2**). However, one of the 26 target genes in each replicate failed to reach the required 50X coverage, a slight reduction in performance compared to the bind60 and lig30 conditions alone (**Supplementary Table 2**, **Figure S4**). The percentage of reads aligned to the target regions for bind60, lig30, and bind60lig30 were 2.8%, 2.7%, and 2.8%, respectively. The combination of extended dA-tailing and ligation steps (bind60lig30) did not lead to a significant improvement in either read number or on-target coverage. In contrast, the bind60 and lig30 conditions produced the most consistent results. Since both bind60 and lig30 demonstrated improvements in sequencing metrics, we ultimately chose the lig30 protocol for routine use, as it is easier to implement in a diagnostic laboratory setting (refer to **Figure 2A** for the finalized library preparation workflow).

**Figure 2.**
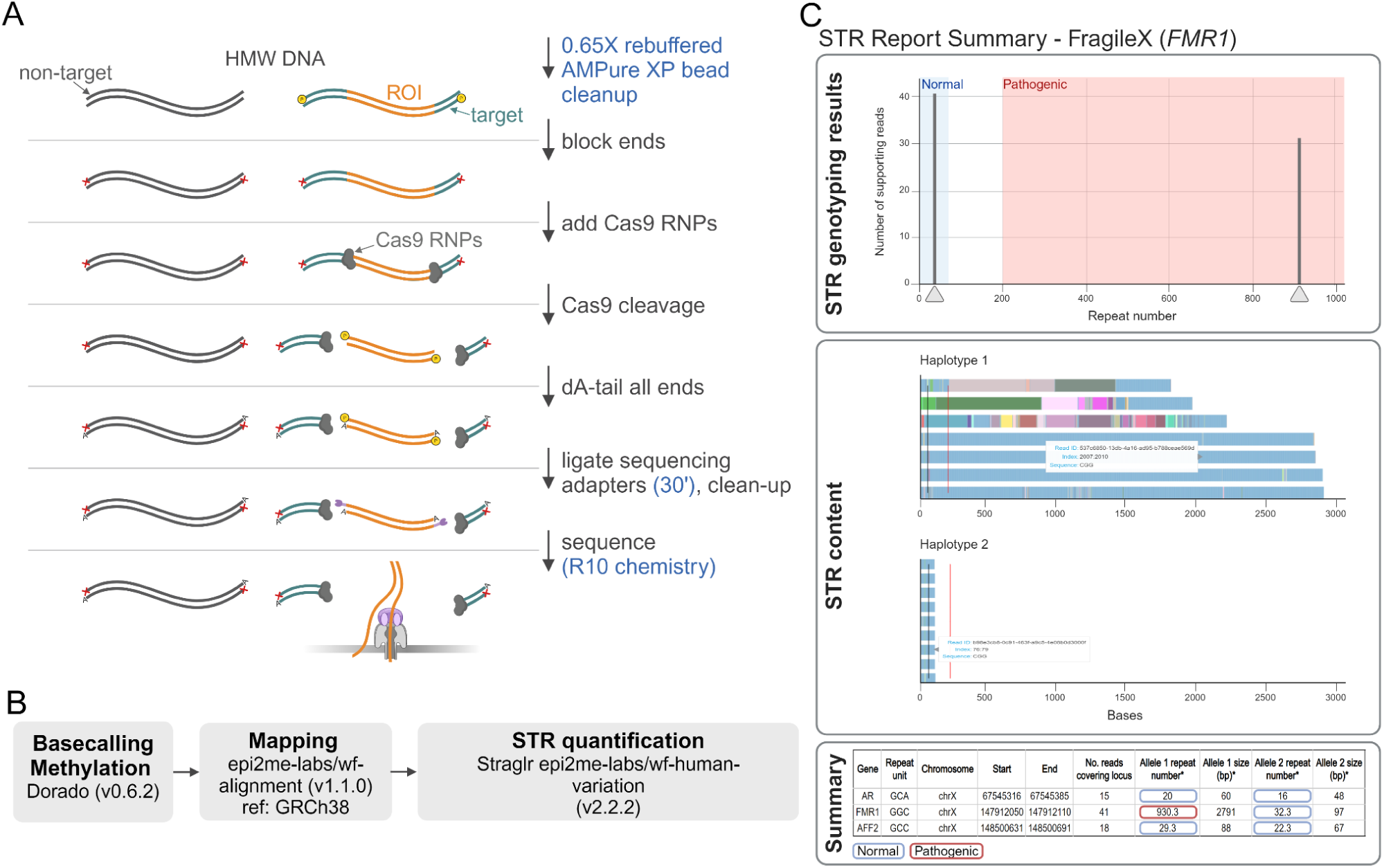
Updated Clin-CATS nanopore Cas9 sequencing workflow for R10 flow cells. **(A)** Library preparation: High-molecular-weight (HMW) DNA extracted from EDTA blood is cleaned up using 0.65X rebuffered AMPure XP beads and then phosphorylated to block the ends, preventing downstream ligation. Cas9 ribonucleoprotein (RNP) complexes are added to excise the region of interest (ROI) (orange) from target DNA (green). All DNA is dA-tailed, and ONT sequencing adapters (purple) are ligated to ROI DNA with a 5’ phosphate and complementary dA tail. The sample is cleaned up and sequenced. Changes specific to the R10 flow cell library preparation protocol are indicated in blue. **(B)** Bioinformatic pipeline: basecalling and methylation are performed with Dorado (v0.6.2), then reads are mapped to the GRCh38 reference genome using epi2me-labs/wf-alignment (v1.1.0), and short tandem repeats (STRs) are quantified using Straglr in the ONT epi2me-labs/wf-human-variation workflow (v2.2.2). **(C)** Example STR report for a FragileX patient: the report includes STR genotyping results showing non-pathogenic (blue) and pathogenic (red) thresholds, repeat configurations for each read, and a summary table with gene, repeat unit, chromosome, positions, number of reads, and allele sizes with their pathogenicity indicated by color.

### Multiplexed R10 Clin-CATS library preparation protocol yields inadequate coverage for diagnostics

Next, we aimed to develop a multiplexed Cas9-based nanopore sequencing protocol to increase throughput and reduce costs for the Clin-CATS workflow. The first step was to optimize the DNA clean-up using rebuffered AMPure XP beads, testing a range of bead ratios from 0.1X to 0.7X to selectively remove short DNA fragments that could reduce sequencing efficiency (**Figure S5**, **Supplementary Table 3**). Our experiments showed that concentrations below 0.5X caused significant DNA loss, while higher concentrations retained more long fragments needed for successful sequencing. Ultimately, a 0.65X bead ratio was identified as optimal, striking a balance by minimizing DNA loss and retaining sufficiently long fragments. Using rebuffered beads, this clean-up strategy successfully increased the DNA Integrity Number (DIN) by an average of 1.2, improving the quality of DNA fragments for nanopore sequencing (**Supplementary Table 3**). Incorporating this method into the routine Clin-CATS diagnostic workflow enabled the recovery of previously failed samples.

Attempts to implement multiplexing using ONT’s beta-testing protocol and several modifications were met with limited success (see **Figure S6** for the library preparation schematic and **Supplementary Table 4** for tested adaptations to the beta-testing protocol ONT multiplexing protocol). We explored multiple approaches, including exonuclease treatments to remove off-target DNA, dephosphorylation to reduce non-specific adapter ligation, and the addition of Proteinase K to remove bound proteins such as Cas9 from the target DNA. Despite these efforts, none of the modifications led to improved performance. Exonuclease-treated samples produced no usable reads, while both Proteinase K and dephosphorylation led to significant reductions in sequencing coverage compared to the unmodified protocol (**Supplementary Table 5, Supplementary Table 6, Figure S7**). Additionally, testing both PAM-in and PAM-out Cas9 guide orientations—where PAM-in refers to the guide RNA being designed with the Protospacer Adjacent Motif (PAM) site facing toward the target region, and PAM-out refers to the PAM site facing away from the target region (see **Figure S8** for a schematic)—did not prevent target DNA from being digested by exonucleases (**Supplementary Table 5**, **Figure S7**). As a result, neither orientation was able to protect the region of interest, leading to poor enrichment and coverage across all tested conditions.

Multiplexed sequencing runs with 2, 4, and 6 samples failed to reach the required 50X coverage, achieving, on average, 3-5X coverage (**Figure 3**, **Supplementary Table 7**, **Supplementary Table 8**). Attempts to increase sequencing yield by reloading flow cells with 2 samples after 24 hours, either once (48h run) or twice (72h run), also resulted in low coverage, with averages of 11.4X and 7.9X, respectively (**Supplementary Table 8**). In subsequent tests, when samples were sequenced individually on R10 flow cells using the barcoding protocol, only two out of six samples achieved 50X coverage for most genes (**Figure S9A**, **Supplementary Table 9**), which was still lower than the coverage obtained in earlier singleplex runs without barcoding on R9 flow cells (**Figure S9B**, **Supplementary Table 9**). The likely cause for these failures is the additional ligation step required to add barcode adapters in the multiplex protocol. Ligation is already inefficient, and introducing another step likely reduces the overall efficiency. This explains why even single samples sequenced with the multiplex protocol failed to reach the required coverage, as the added barcoding step further compromised the library preparation.

**Figure 3.**
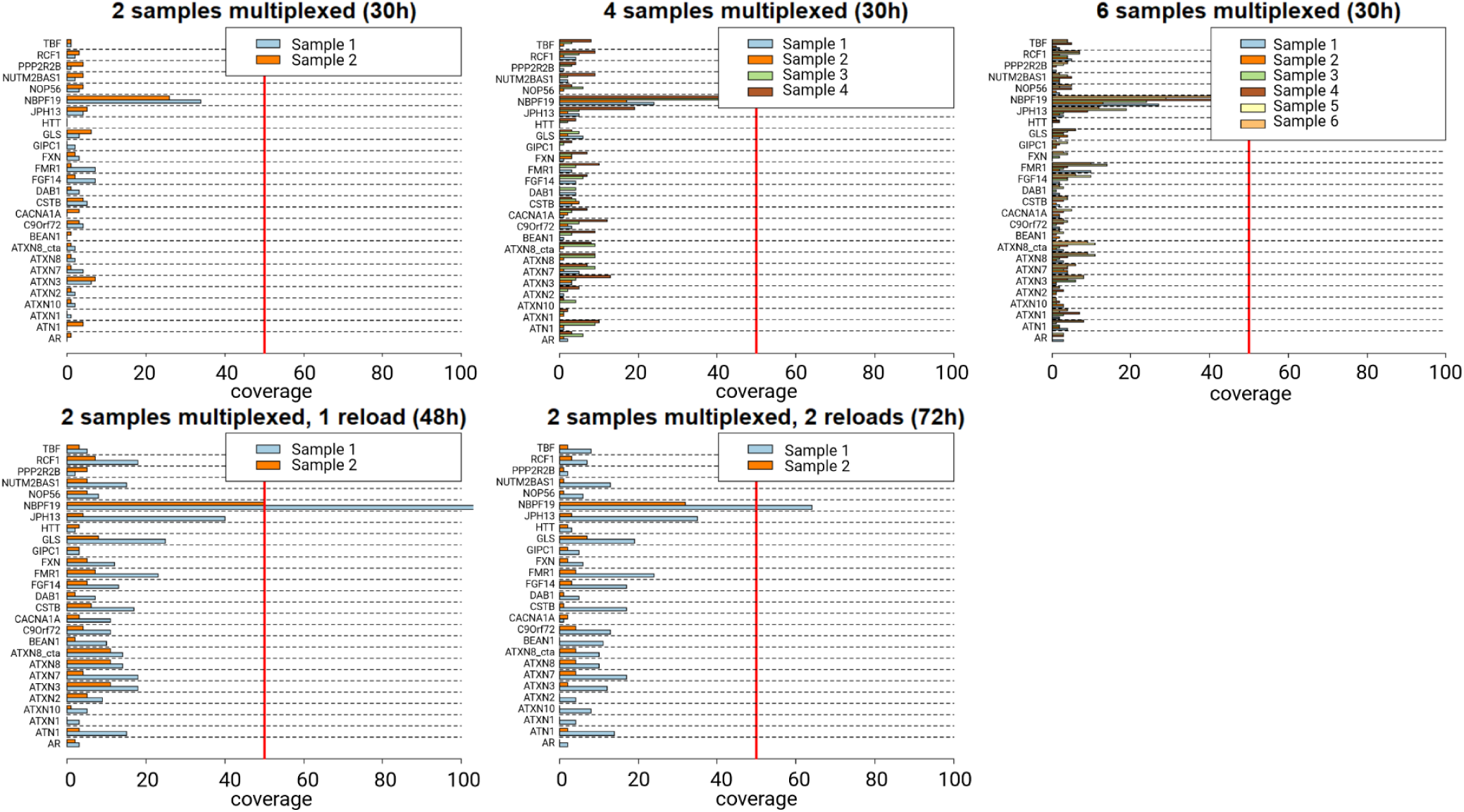
Coverage for multiplexing tests with the ataxia panel. Boxplots of sequencing coverage for multiplexing tests with 2, 4, and 6 samples over a 30h runtime. Coverage for 2 multiplexed samples with a 48h runtime, including one library recovery and reloading at 24h. Coverage for 2 multiplexed samples with a 72h runtime, including two rounds of library recovery and reloading at 24h and 48h. The target coverage of 50X is indicated by a red line (n=2-6 biological replicates as indicated in the figure legend). Refer to **Supplementary Table 7** for read counts and **Supplementary Table 8** for quantification.

### Integration of Straglr with expanded gene set into the Clin-CATS workflow ensures R10 compatibility

In our effort to update the Clin-CATS bioinformatics workflow for the new R10 flow cells, we considered several tools for analyzing repeat expansions. The R10 flow cells, which are set to replace the currently used R9 flow cells, are not yet supported by the STRique tool (as noted in <u>STRique’s GitHub repository</u>). This lack of support, coupled with the need to re-train the pore model to adapt to the R10 chemistry, made STRique less viable for our updated pipeline. Additionally, STRique is incompatible with ONT’s new POD5 data format, and its development has been largely stagnant, leaving little room for future improvements or troubleshooting. As a result, we opted to explore alternative tools that could better handle the specific requirements of R10 flow cell data. Tools such as Straglr, tandem-genotypes^16^, TriCoLoR^17^, and RepeatHMM^18^ were considered. Based on recommendations from ONT and the nanopore community, its integration into the ONT Epi2Me wf-human-variation, as well as the fact that other tools either lacked support for newer algorithms or required complex adjustments, Straglr, in particular, stood out as a promising candidate. Moreover, Straglr was selected for integration into the Clin-CATS pipeline due to its ability to calculate repeat units post-basecalling, which increases accuracy by mitigating sequencing errors and biases and allows it to be used for future flow cell chemistry changes. In the updated Clin-CATS workflow, basecalling and methylation calling are performed with Dorado (v0.6.2), reads are then aligned to the GRCh38 reference genome using epi2me-labs/wf-alignment (v1.1.2) and STRs are quantified using Straglr within the ONT epi2me-labs/wf-human-variation workflow (v2.2.2) (**Figure 2B**). Furthermore, ONT has incorporated Straglr into an Epi2Me Nextflow workflow, offering a user-friendly HTML report with a clear visualization of repeat expansions (**Figure 2C**).

We adjusted the Straglr pipeline from the ONT repository to include missing genes from our panels (**Supplementary Table 10**). Several genes were already present in both the Straglr pipeline and Clin-CATS, including *AR*, *ATN1*, *ATXN1*, *ATXN10*, *ATXN2*, *ATXN3*, *ATXN7*, *ATXN8OS/ATXN8*, *C9orf72*, *CACNA1A*, *FMR1*, *FXN*, *GIPC1*, *HTT*, *JPH3*, *LRP12*, *NOP56*, *NOTCH2NLC/NBPF19*, *PPP2R2B*, and *TBP*. However, several key genes from the Clin-CATS panel were missing from the Straglr pipeline, including *BEAN1*, *DAB1*, *FGF14*, *LOC642361/NUTM2B-AS1*, *RFC1*, and *RILPL1*, which we subsequently added. This ensured the comprehensive coverage of clinically relevant genes across both the Clin-CATS and Straglr workflows.

### Validation of the updated Clin-CATS workflow shows improved accuracy but reduced coverage

We validated the updated Clin-CATS workflow, which incorporates the adapted library preparation protocol, sequencing on R10 flow cells, and analysis using Straglr in the ONT wf-human-variation Epi2Me pipeline (as shown in **Figure 2**). We compared this updated workflow to the original Clin-CATS setup, which used the original Cas9 library preparation, R9 flow cells, and STRique analysis (per Erdmann et al. 2023^8^). This evaluation used clinical samples to assess both workflows’ accuracy in detecting known repeat unit (RU) sizes across genes in the Ataxia panel (**Supplementary Table 11**). Samples with high coverage from the original workflow were selected for this comparison.

Both workflows produced consistent results for *ATN1*, detecting 12/50 RUs, and for *ATXN2*, accurately identifying the pathogenic allele of 32 RUs. For *ATXN1*, the workflows closely matched: the original workflow reported 32/49 RUs, while the updated reported 32/48 RUs. For *ATXN3*, the original workflow identified 72 RUs, while the updated workflow was closer to the true repeat expansion size, overestimating by 1 repeat unit. For *ATXN8*, the updated workflow reported a median RU size for the pathogenic allele closer to the expected median.

The original workflow with STRique could not detect repeat expansions in *BEAN1*, and the two alleles were quantified manually in IGV. In contrast, the updated Clin-CATS workflow successfully identified non-pathogenic TAAAA repeat expansions of 18/857 and 19/640 RUs in two SCA31 patients. For a patient with a repeat expansion range of 381-1895 RUs and a median 927 RU in *C9orf72*, the original workflow reported 4/7-2697 GCCCCG repeats, whereas the updated workflow reported 4/556 RUs. The results for *CACNA1A* were consistent in both workflows, with both detecting 13/23 repeats, close to the expected number of RUs. For *FGF14*, both workflows agreed within the pathogenic range of 252-312 RUs, though the updated workflow offered a median value of 290 RUs, and the original a range of 60-372 RUs in the pathogenic expansion. For *FMR1*, the updated workflow accurately identified 19/56 RUs for both alleles, while the original reported a broader range (21/27-77 RUs). For *FXN*, the updated workflow yielded a more precise 104/1093 RU result compared to the original workflow’s broader range of 63-1569 RUs, both within the pathogenic range of 100/850-950 RUs. For *RFC1*, the original workflow provided incorrect allele counts, while the updated workflow accurately detected the correct RU count, though it slightly overestimated the pathogenic allele repeat count.

The R10 flow cells provided improved mean read quality (16.5) over R9 (15.3), suggesting enhanced sequencing accuracy across clinical samples (**Supplementary Table 11**). However, the updated workflow, when applied to the same samples used in the original workflow, produced lower coverage. Mean coverage across all targets for the original workflow was 439X, compared to 94X in the updated workflow. Target-specific coverage was similarly lower, averaging 444X for the original versus 75X for the updated. This coverage reduction can be attributed partly to sample age (2-3 years) and the impact of freeze-thaw cycles. To better assess the updated Clin-CATS workflow’s coverage, freshly collected and extracted control samples were sequenced, yielding an average coverage of 222X across all targets (**Supplementary Table 12**), which remains approximately 40% below that achieved with the original workflow. Additionally, Coriell samples processed with the updated workflow achieved a read quality of 17.6 but displayed lower-than-expected coverage, averaging 185X across all targets and 125X for specific target genes (**Supplementary Table 13**).

### The Clin-CATS workflow on R10 flow cells accurately detects repeat expansions in Coriell samples

We further validated the updated Clin-CATS workflow using Coriell samples (**Supplementary Table 13**), focusing on detecting known repeat expansions in genes associated with various neurological conditions. In the SBMA sample NA23709, the *AR* CAG repeat was correctly identified with a median count of 49/51 RUs. In DRPLA samples NA13716 and NA13717, expansions in the *ATN1* gene were detected with median counts of 20/72 RUs and 18/72 RUs, respectively, matching the expected ranges. For DM1, sample NA03759 showed an expanded CTG repeat in *DMPK* with a median count of 15/2317 RUs, while NA03990 displayed a repeat range of 15/79 RUs, both consistent with the expected repeat number range. Similarly, the *FMR1* repeat expansion in FXS sample NA09237, with a known range of 931-940 RUs, was detected with a median repeat count of 864/1554 RUs, while samples NA20236 and NA20239 also matched the expected repeat number ranges for normal and premutation statuses, respectively. For FRDA, sample NA16243 with an *FXN* expansion had a median GAA repeat count of 515/1953 RUs, fitting within the expected double expansion range. In the case of HD, we could detect the correct number of *HTT* CAG repeats in samples NA13505 (17/45), NA13505 (22/50), NA13507 (15/55), NA13508 (22/58), NA13513 (15/49), NA13514 (15/52) and NA13512 (16/44), while NA13503 and NA13515 matched the expected range.

## DISCUSSION

In this study, we presented an updated Clin-CATS workflow compatible with ONT R10 flow cells, expanding the panel to 27 genes associated with ataxia, myopathy, neurodegeneration, and ALS/motor neuron diseases. This update brings several advantages in terms of diagnostic utility, sequencing accuracy, and workflow flexibility but also highlights significant challenges, particularly in the context of multiplexing and coverage.

The expanded gene panels significantly increase the diagnostic potential of Clin-CATS by enabling the simultaneous investigation of multiple repeat expansion-related genes across various neurological conditions (**Figure 1**, **Supplementary Table 1**). Another necessary refinement in the updated Clin-CATS workflow was the modification of the Cas9 library preparation protocol to address the suboptimal performance of the Cas9 approach on R10 flow cells. Although ONT has discontinued their Cas9 protocol, equivalent reagents, such as tracrRNAs and Cas9 Nuclease, can be purchased from IDT and NEB, allowing the continuation of the workflow (refer to *Methods* sections *Library preparation and ONT Cas9-targeted sequencing*). The final optimized protocol includes cleaning the samples with 0.65X rebuffered AMPure XP beads to remove short DNA fragments and extending the adapter ligation time from 10 to 30 minutes (**Figure 2A**). These modifications were essential for improving sequencing efficiency and ensuring that the R10 flow cells could meet the required 50X coverage thresholds for clinical diagnostics (**Supplementary Table 2**).

In this study, we attempted to multiplex samples prepared for Cas9 R10 sequencing by experimenting with several key parameters. With only 2.7% of reads aligning to the target region on average (**Table 3**), we explored methods to reduce off-target DNA and improve dA tailing and ligation efficiency by freeing the 3’P ends of the on-target DNA. Key adaptations included optimizing guide RNA orientation to control Cas9 binding specificity (on-target or off-target), utilizing Proteinase K to digest Cas9 proteins that purportedly adhere to the template post-cleavage to free the DNA ends for dA tailing and ligation, applying a combination of exonucleases to digest off-target DNA, and employing dephosphorylation to differentiate the 5’ ends of the target DNA from off-target DNA, which retains a 5’P after Cas9 cleavage. Despite these efforts, multiplexing failed to achieve the necessary coverage for clinical diagnostics. Sequencing coverage for multiplexed samples fell below the required 50X threshold, with average coverage ranging from 3-5X, even when multiple samples were reloaded onto flow cells and the sequencing run time was extended (**Figure 3**, **Supplementary Table 8**). While previous studies have demonstrated successful multiplexing of Cas9-enriched libraries^19,20^, they typically targeted fewer genes, resulting in higher sequencing efficiency. In contrast, our larger gene panel exceeded the current capabilities of multiplexed Cas9-enriched nanopore sequencing, particularly with R10 flow cells. The challenges of multiplexing in this study likely stemmed from the inefficiencies introduced by additional ligation steps required for barcoding each sample, which may have further reduced overall library preparation efficiency. To overcome these limitations, protocol refinements are needed, such as exploring higher-throughput sequencing platforms like PromethION, which may be better suited for multiplexed workflows involving extensive gene panels. Additionally, a critical discovery in earlier studies indicated the necessity of using fresh blood samples for DNA extraction. Current DNA extraction via an automated system does not optimize for high-molecular-weight (HMW) DNA, which, while practical for diagnostics, could benefit from alternative methods like Nanobind to further enhance coverage and sequencing performance.

While R10 flow cells deliver substantial improvements in sequencing accuracy, the overall coverage achieved with R10 was lower compared to that of R9 flow cells. Specifically, our results showed that the updated workflow, which employs R10 flow cells, achieved a mean coverage of 222X across all targets (**Supplementary Table 12**), compared to 439X for the original workflow, which uses R9 flow cells (**Supplementary Table 11**). This lower coverage could present a challenge for Cas9-targeted sequencing applications, which require consistent coverage across target regions. Despite this limitation, the updated Clin-CATs approach met the 50X minimum coverage threshold required for our clinical diagnostics workflow. Moreover, the updated workflow demonstrated improved accuracy in detecting and quantifying repeat units in both clinical and Coriell samples (**Supplementary Table 11**, **Supplementary Table 13**). It is worth noting that our updated Cas9-targeted sequencing platform using R10 flow cells provides greater coverage than what is typically achieved with adaptive sampling, reinforcing its advantage for applications requiring higher sequencing depth and accuracy.

R10 flow cells present clear benefits, offering higher accuracy and, in non-Cas9-applications, increased throughput. As ONT continues refining their technology, we anticipate even greater throughput from R10 flow cells across various protocols, including those based on Cas9. Although adopting ONT’s Q line—providing guaranteed R9 flow cell availability for a fixed period—could support maintaining higher coverage, this option restricts the potential to incorporate future upgrades and reduces operational flexibility by committing the sequencing device exclusively to R9 flow cells. Given ONT’s rapid advancements, we believe transitioning to R10 flow cells is a more strategic long-term solution.

An important improvement in the updated Clin-CATS workflow was the integration of Straglr^12^, a tool embedded in the ONT epi2me-labs/wf-human-variation workflow, which processes basecalled reads rather than raw electrical signals. The original Clin-CATS workflow relied on the now-deprecated tool STRique^7^, which quantified STRs from raw signal data. Our original Clin-CATS pipeline using STRique was generally incapable of determining a median for large repeat expansions (>100 RU) and instead provided a range of repeat sizes; as a result, these expansions, for example, in the case of *RFC1*, required manual evaluation. The rationale for this approach is that large expansions exhibit a broad range of repeat sizes, leading to a lack of a single peak for one repeat size, as seen with shorter expansions. Following distribution plotting, peak determination was conducted manually, though retrospectively, this approach introduced limitations in precision. Moreover, the migration to Straglr enhances Clin-CATS compatibility with future ONT flow cell chemistries, ensuring long-term adaptability. By processing basecalled data, the pipeline can function seamlessly across different sequencing chemistries, reducing the need for revalidation or retraining models, an issue that was more prevalent with tools like STRique, which were tied to specific pore models and data formats. Additionally, using ONT’s workflow ensures that the pipeline remains actively maintained.

While Straglr presents a robust advancement in the Clin-CATS workflow, some gene-specific adjustments are necessary to ensure optimal performance. Challenges persist in achieving precise phasing and accurately handling subtle repeat differences, especially in differentiating closely sized alleles, such as a 13-repeat versus a 14-repeat allele. Additionally, we used Straglr parameters that do not consider broken reads, which may limit its effectiveness in capturing long repeats, especially for genes like *BEAN1*, *C9orf72*, and *RFC1*, where pathogenic repeat expansions are extensive. Importantly, for the diagnosis of these conditions, it is often sufficient to detect an expansion exceeding a specific threshold, an assessment that can be made even from broken reads. This threshold-based detection approach may also explain the divergent results observed for *C9orf72*. Improvements in DNA extraction and library preparation could help preserve DNA fragments, increasing the likelihood of full-length reads and improving accuracy for these complex loci. However, expansions of the pathogenic AAGGG repeat motif in *RFC1* seem to be prone to breaking or fragmenting, likely due to the formation of specific secondary DNA structures^21^, presenting an inherent challenge beyond library preparation alone. Additionally, the *FMR1* repeat expansion in the FXS sample NA09237, with a known range of 931-940 RUs, was detected with a median repeat count of 864/1554 RUs. This discrepancy likely stems from the sample being male, containing only one FMR1 copy, which Straglr did not account for, instead forcing a two-allele call. Future refinements could incorporate gender-specific handling in X-linked disorders to improve accuracy.

The reporting approach of Straglr could be further improved, particularly for genes like *RFC1* that contain multiple repeat motifs. Currently, extracting details about repeat interruptions or expansions involving multiple motifs requires manually examining individual reads, which is not ideal for clinical reporting. Enhancing Straglr’s ability to consolidate and report complex repeat structures directly in summary outputs would significantly improve its clinical utility, particularly for diagnosing disorders with heterogeneous repeat motifs.

STRique does not further process the isolated repeat lengths of individual reads and lacks phasing capabilities, introducing ambiguity that complicates precise allele differentiation and often necessitating intensive additional inspection of reads in certain cases. In contrast, Straglr reports the median repeat count, offering a more consistent and interpretable result that supports clearer allele identification. Straglr also provides key advantages, including compatibility with evolving ONT technologies, enhanced bioinformatics support, and active development, making it the preferred choice for the Clin-CATS workflow. This shift not only improves the pipeline’s adaptability and precision but also ensures alignment with ONT’s latest sequencing platforms, positioning the workflow for future advancements in clinical diagnostics.

In conclusion, the updated Clin-CATS workflow, which incorporates an adapted library preparation protocol, R10 flow cell sequencing, and analysis with Straglr in the ONT wf-human-variation Epi2Me pipeline, ensures continuity of this approach despite the discontinuation of R9 flow cells and the Cas9 library preparation kit, while also providing substantial benefits. However, challenges such as suboptimal multiplexing and reduced sequencing coverage underscore the need for ongoing refinements. Despite these hurdles, the workflow demonstrates strong adaptability with expanded panels, improved sequencing accuracy, and bioinformatic flexibility, supporting its long-term application in research and clinical diagnostics. Future efforts should focus on optimizing multiplexing protocols, increasing throughput to fully leverage R10 flow cell capabilities, improving library preparation protocols to preserve better full-length reads for accurate large expansion analysis, and enhancing bioinformatic tools like Straglr for better reporting of complex repeat expansions.

## SUPPLEMENTAL FIGURES

**Figure S1.**
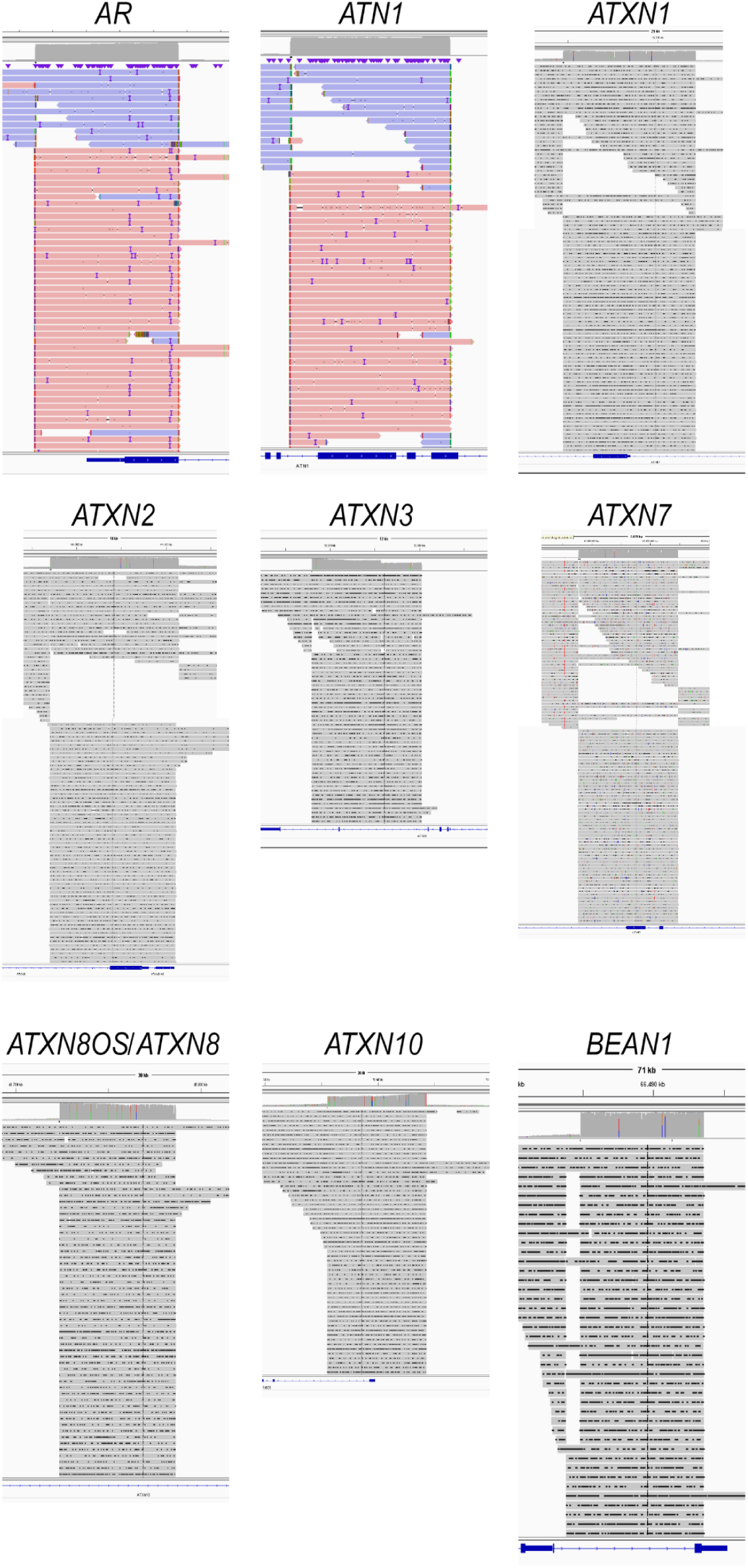
IGV snapshots confirming guide RNA targeting of the correct loci for *AR*, *ATN1*, *ATXN1*, *ATXN2*, *ATXN3*, *ATXN7*, *ATXN8OS/ATXN8*, *ATXN10*, and *BEAN1* genes. Detailed information regarding the specific target regions and guide RNAs can be found in **Supplementary Table 1**.

**Figure S2.**
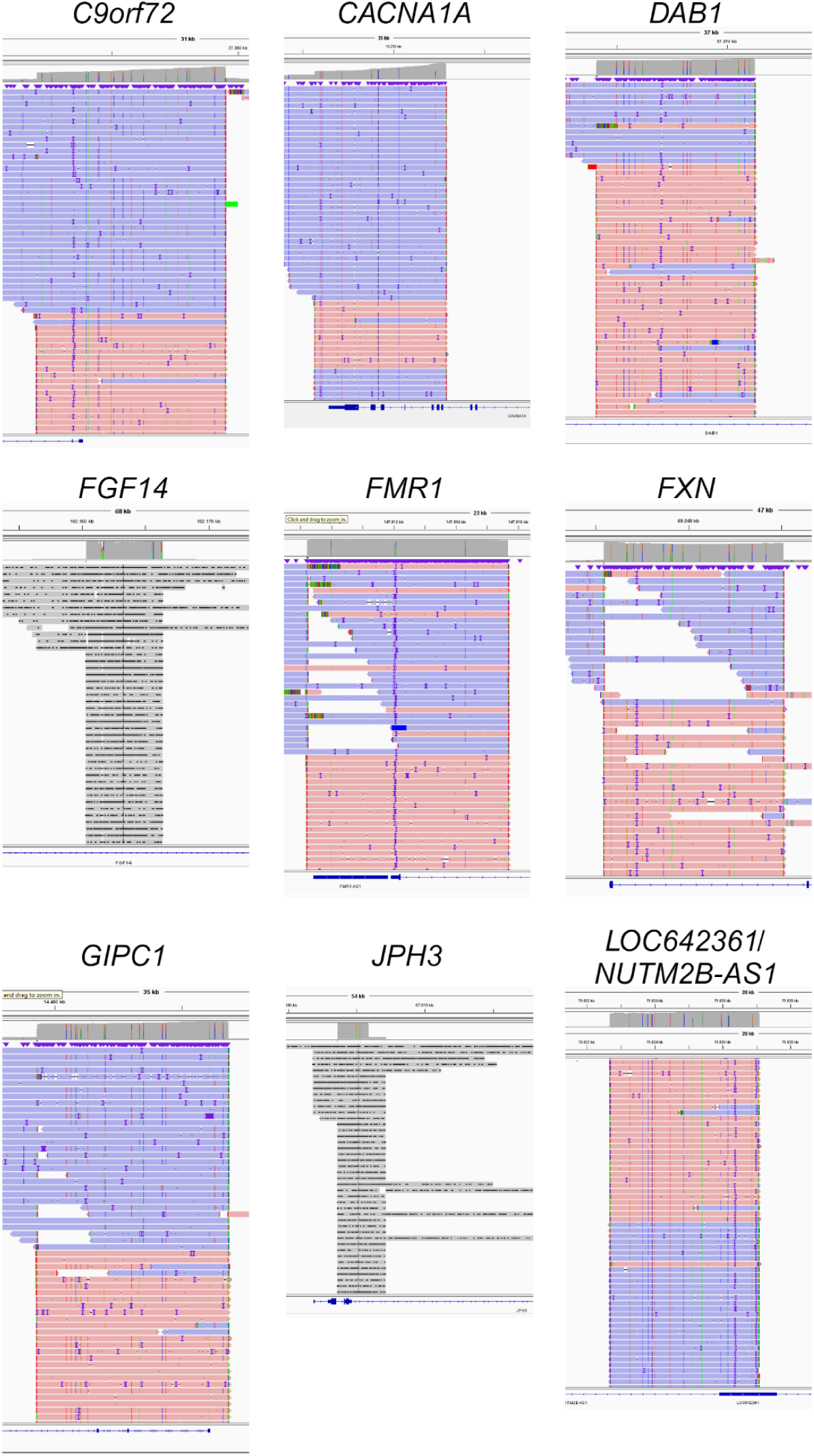
IGV snapshots confirming guide RNA targeting of the correct loci for *C9orf72*, *CACNA1A*, *DAB1*, *FGF14*, *FMR1*, *FXN*, *GIPC1*, *JPH3*, and *LOC642361/NUTM2B-AS1* genes. Detailed information regarding the specific target regions and guide RNAs can be found in **Supplementary Table 1**.

**Figure S3.**
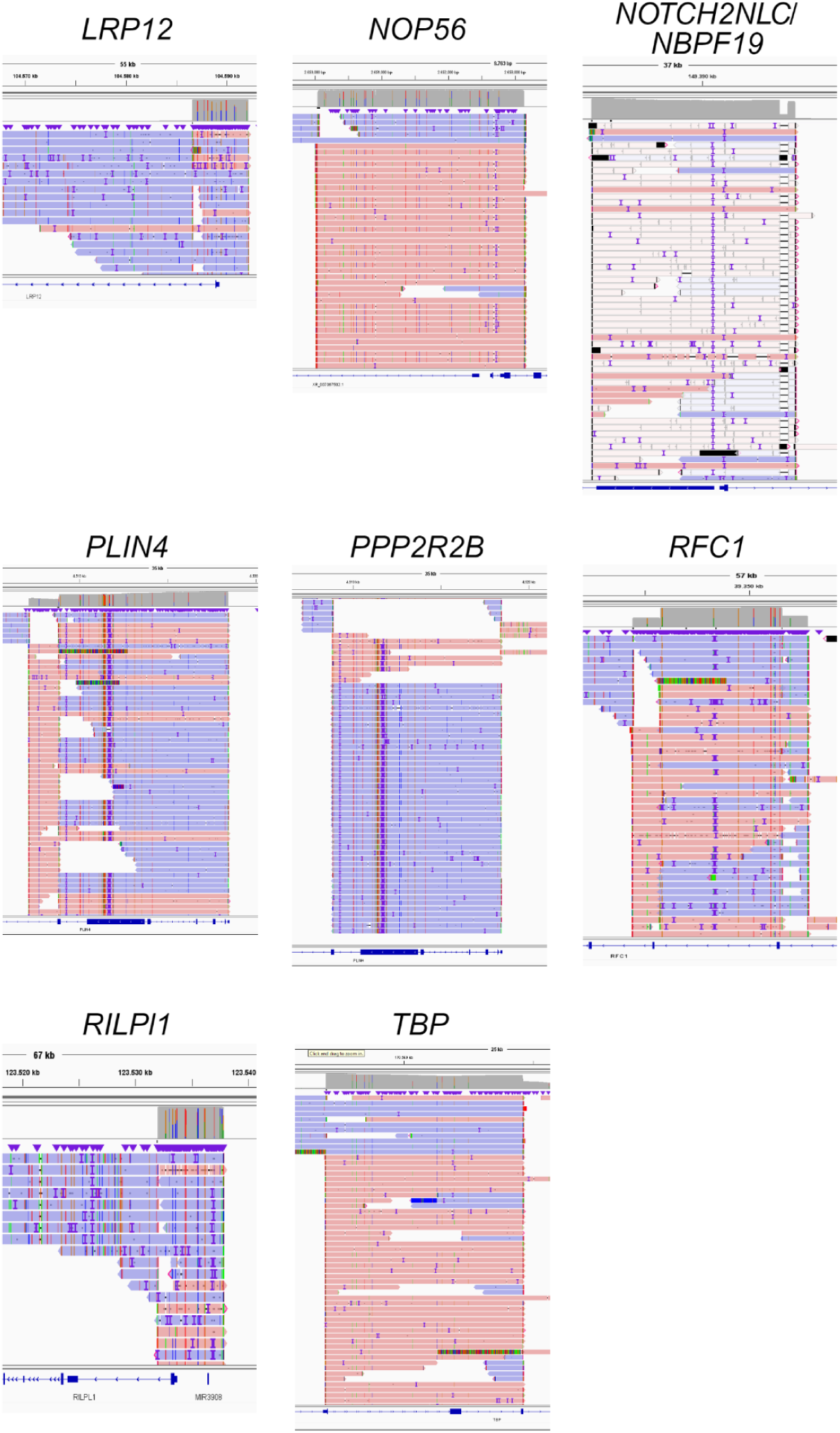
IGV snapshots confirming guide RNA targeting of the correct loci for *LRP12*, *NOP56*, *NOTCH2NLC/NBPF19*, *PLIN4*, *PPP2R2B*, *RFC1*, *RILPL1*, and *TBP* genes. Detailed information regarding the specific target regions and guide RNAs can be found in **Supplementary Table 1**.

**Figure S4.**
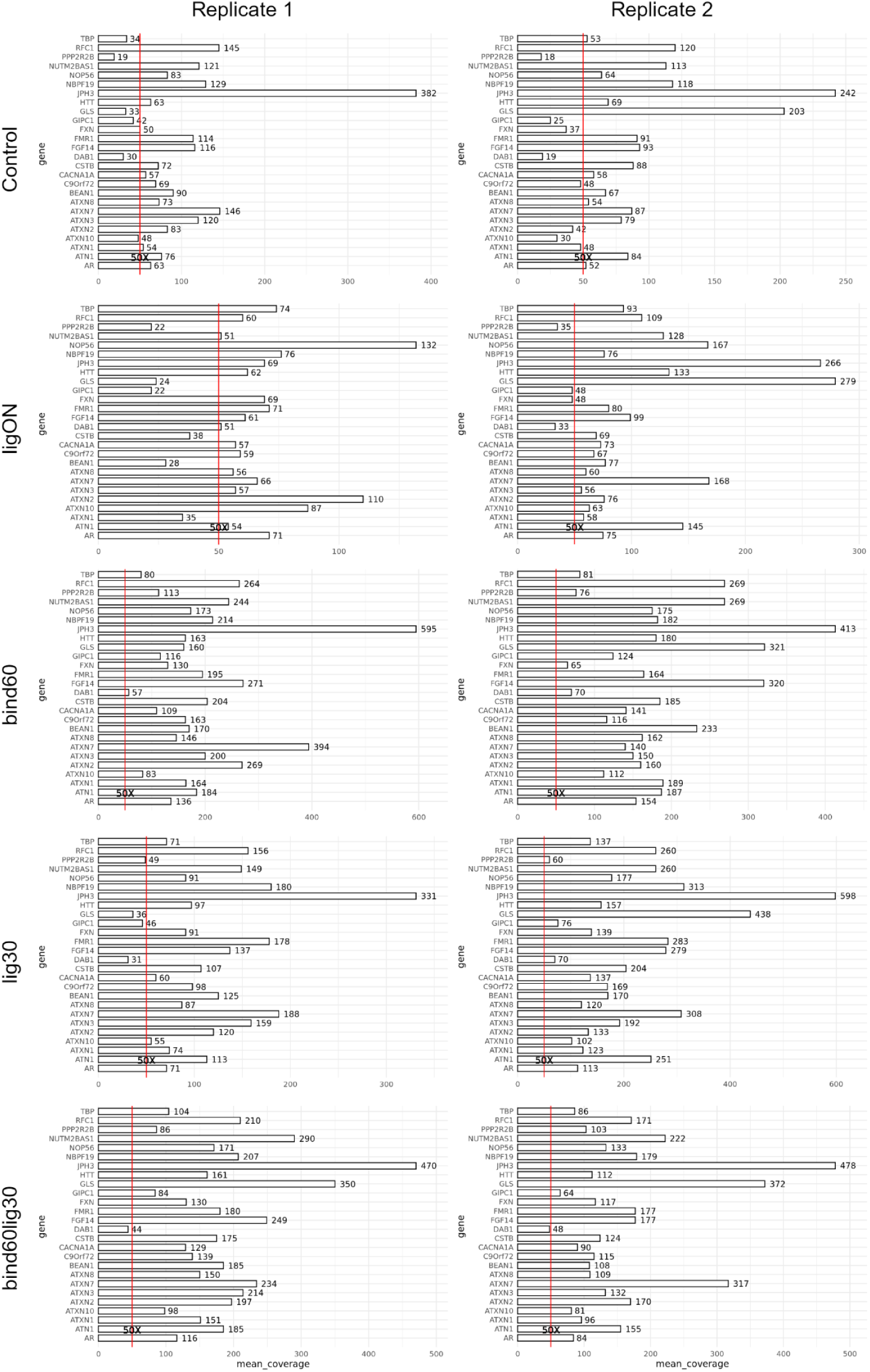
Coverage across target regions of the Clin-CATS Ataxia panel for adaptations of the Cas9 protocol sequenced on R10 flow cells. Barplots showing the mean coverage for the repeat target region of each gene in the Ataxia panel for n=2 technical replicates. The red line represents the 50X coverage threshold. Numbers to the right of each bar indicate the mean coverage for the target gene for that specific run. The control condition is the original Cas9 protocol sequenced on R10 flow cells.

**Figure S5.**
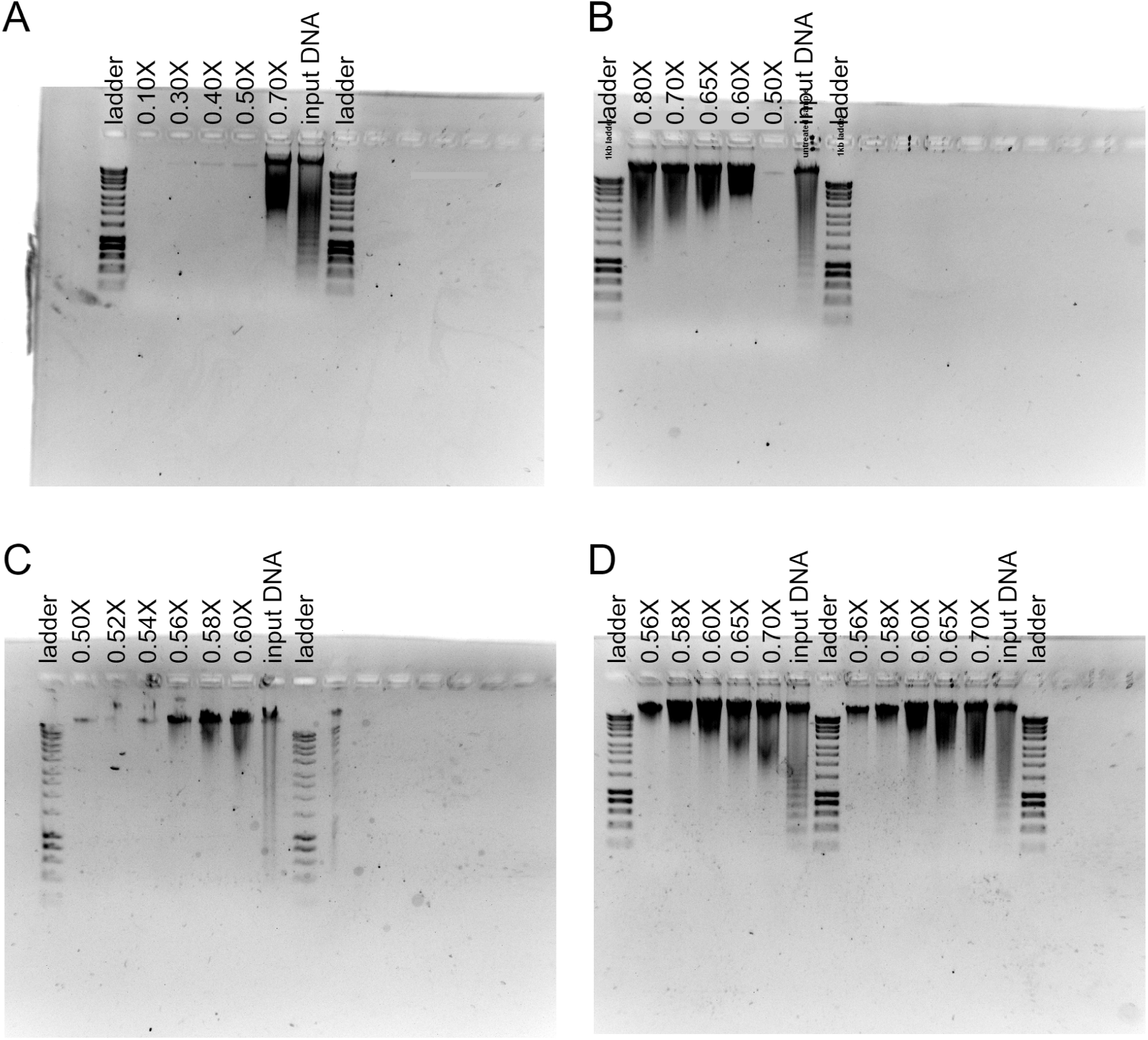
Agarose gels showing the results of AMPure XP bead clean-up tests for input DNA used in Cas9 library preparation. **(A)** Initial clean-up test with 0.1X-0.7X, **(C)** 0.8X-0.5X (Test 1), **(C)** 0.6X-0.5X (Test 2), and **(D)** two biological replicates of 0.7X-0.56X (Test 4.1-4.2). Refer to **Supplementary Table 3** for the DNA quantification from Tests 1-4.2.

**Figure S6.**
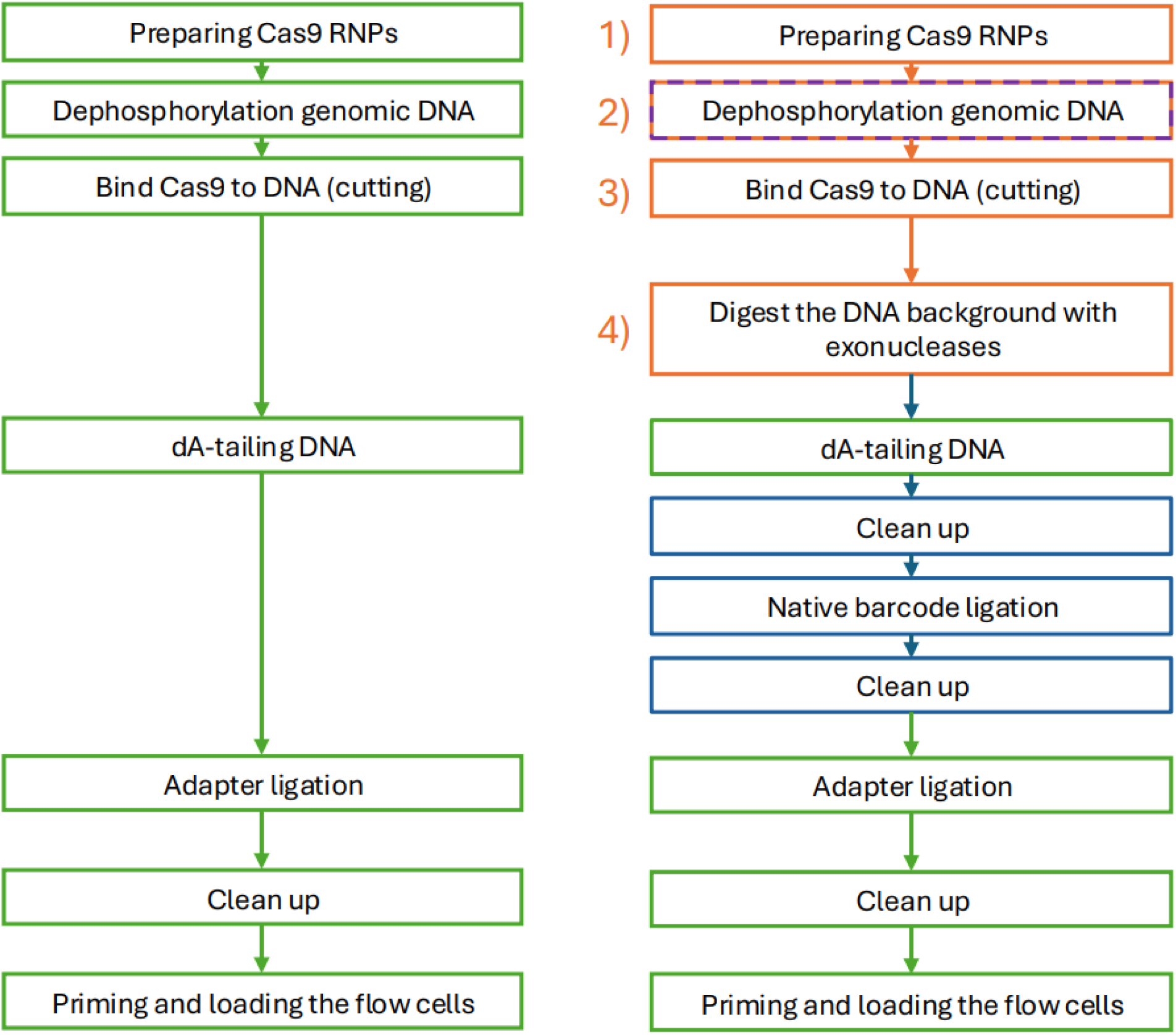
Flow chart showing the workflow and changes of the multiplex approach. Numbers indicate the protocol adaptations tested: 1) PAM-in or PAM-out orientation guides, influencing whether the Cas9-protein sticks to the target or off-target DNA. 2) DNA dephosphorylated to improve the efficiency of dA-tailing on the target DNA. 3) Proteinase K is used to remove the remaining Cas9 protein that may be adhered to the region of interest (ROI). 4) Digestion of off-target DNA using exonucleases to increase the relative percentage of target DNA.

**Figure S7.**
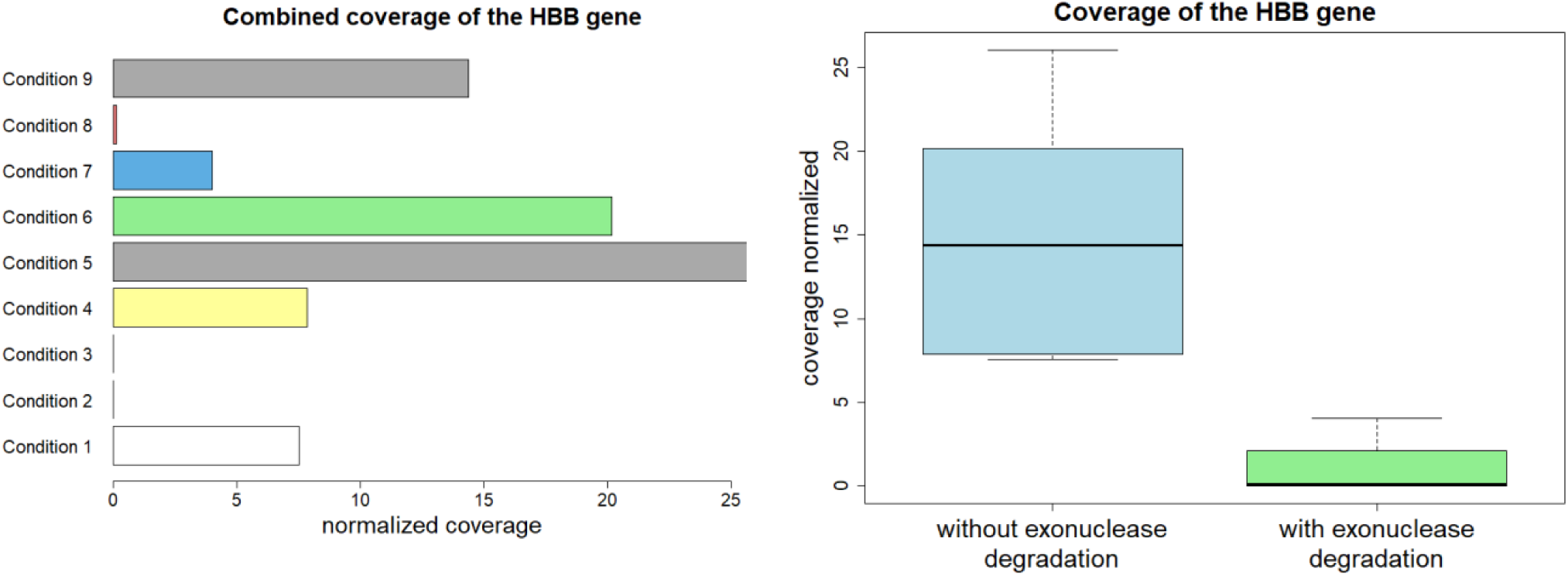
Effects of multiplexing protocol adaptations on sequencing coverage. Barplot of the mean coverage of each library preparation condition, normalized to pore count (n=2 biological replicates) and boxplot comparing the coverage normalized to pore count with and without exonuclease degradation (n=2 biological replicates). Refer to **Supplementary Table 5** for the relevant data.

**Figure S8.**
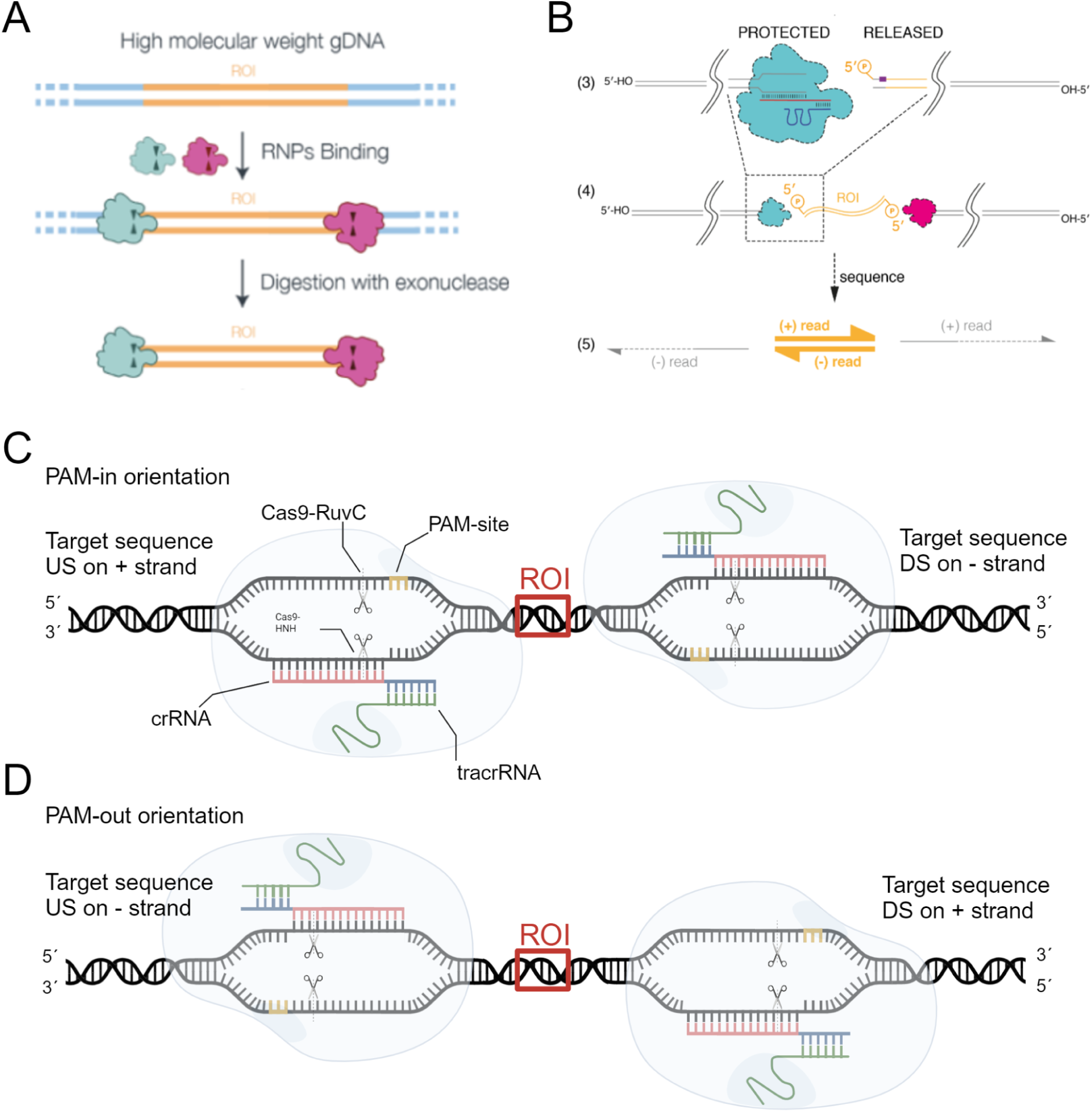
Orientation of the Cas9 protein after cleavage. **(A)** Schematic of the most recent beta-testing ONT multiplex protocol showing the binding of the Cas9 protein to the target DNA (copied from ONT protocol) **(B)** Schematic of the ONT simplex protocol, showing the Cas9 adhering to the off-target DNA (copied from ONT protocol). **(C)** Schematic of the PAM-in orientation guide RNA design, where the PAM sites are orientated towards each other, and the crRNA sits on the non-target strand. **(D)** Schematic of the PAM-out orientation guide RNA design, where the PAM sites are orientated away from each other, and the crRNA sits on the target strand. RNP ribonucleoprotein; US upstream; DS downstream; ROI Region of Interest; PAM Protospacer Adjacent Motif.

**Figure S9.**
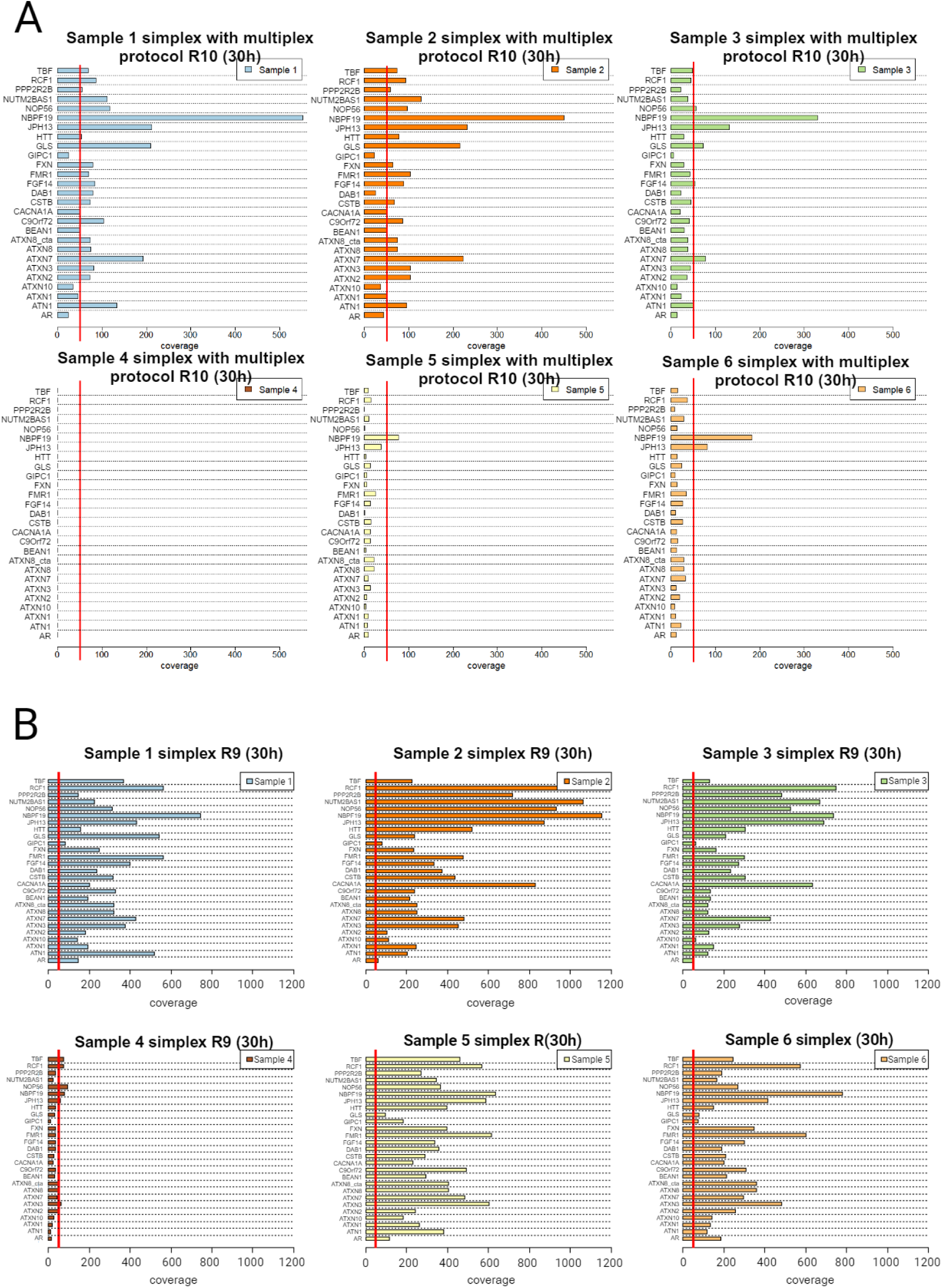
Coverage across target regions of the Clin-CATS Ataxia panel for singleplexed samples. Barplots showing the mean coverage for the repeat target region of each gene in the Ataxia panel for individual samples prepared using the **(A)** multiplexing protocol and sequenced on R10 flow cells and **(B)** original R9 protocol (non-multiplexed) and sequenced on R9 flow cells. The red line represents the 50X coverage threshold. For read counts, refer to **Supplementary Table 9**.

## METHODS

### Guide RNA design

New guides not included in the original Erdmann and Scharf et al.^8^ publication were designed using CHOPCHOP v3.0.0. All guides were designed using the hg38 *Homo sapiens* reference and have a length of 20 bp. The resulting ROI flanked by the guide RNAs should be 3 to 12 kb. Multiple guides up- and downstream with the highest efficiency that were not self-complementary or complementary to any other site by a minimum of 2 mismatches in the list provided by CHOPCHOP were selected. The strand location of the upstream (and downstream) guide was on either the plus or minus strand but always opposite of the other crRNA, depending on the orientation of the Cas9 needed. The most efficient pair was selected, while pairs with a guide below 45% efficiency were discarded. The guide RNA sequences for Clin-CATs are provided in **Supplementary Table 1**. For multiplexing tests, specific *HBB* guide RNAs were selected to target a 400-600 bp region of the HBB gene, ensuring efficient enrichment of fragments within this size range. As before, these guides targeted both upstream and downstream of the desired regions. The *HBB* guide RNA sequences selected were: PAMin upstream with the sequence TTATGTTTCACATATGAGCA, PAMin downstream with the sequence GAAGAATTTTATTATATGAG, PAMout upstream with the sequence AAACTGGGGGATATTATGAA, and PAMout downstream with the sequence TTCTTTAGAATGGTGCAAAG. All guide RNAs were purchased from IDT.

### Patient samples and study approval

Informed consent to participate in this study was obtained from all patients and was approved by local institutions (Bayerische Landesärztekammer, 2019– 210). All genetic analyses and investigations were performed in accordance with the guidelines of the Declaration of Helsinki.

### Extraction of genomic DNA

Genomic DNA (gDNA) was obtained from total peripheral EDTA blood samples by extraction of white blood cells with a Biomek FX or i7system (Beckman Coulter) using the NucleoMag® Blood 3 ml Kit (Machery-Nagel, 744502.1) as per the manufacturer’s instructions. The DNA concentration was measured using a Qubit 4 fluorometer (Thermo Fisher Scientific, Q33238) and the Qubit dsDNA HS Assay-Kit (Thermo Fisher Scientific, Q33233) or Qubit dsDNA BR Assay-Kit (Thermo Fisher Scientific, Q32853) as per manufacturer’s instructions. DNA purity (A260/A280 and A260/A230) was measured with the BigLunatic device (Unchained labs) as per manufacturer’s instructions. The DNA Integrity Number (DIN) was measured on the TapeStation 4200 (Agilent Technologies, G2991BA) using the Genomic DNA Screen Tape (Agilent Technologies, 5067-5365) and the Genomic DNA Reagents (Agilent Technologies, Inc, 5067-5366) as per manufacturer’s instructions.

### DNA clean-up using rebuffed Ampure XP beads

2 mL AMPure XP beads (Beckman Coulter, A63882) were mixed thoroughly by vortexing, pelleted on a DynaMag™-2 Magnet magnetic rack (Thermo Fisher Scientific, 12321D) and the supernatant discarded. The beads were washed with nuclease-free water, resuspended by vortexing, repelleted, and the supernatant removed. The washing step was repeated, and the beads were air-dried. A custom buffer consisting of 10 mM Tris-HCL (Thermo Fisher Scientific, J22638.AE), 1 mM EDTA pH 8.0 (Merck, 324506), 1.6 M NaCl (Merck, 59222C), and 11% PEG 8000 (Sigma Aldrich, SIAL-P1458-25ML)^22^ was added to the beads and mixed by vortexing. The rebuffered Ampure XP beads were stored at 4°C until further use for up to 1 week.

DNA samples were diluted to a minimum of 40 ng/µL and cleaned up using 0.65X rebuffered beads, mixed by flicking, placed into a Thermomixer (Eppendorf, 5382000015) at a speed of 450 rpm, with manual inversion every 2 minutes, for 10 minutes. The beads were pelleted, the supernatant was discarded, and the samples were washed with 200 µL of freshly prepared 70% ethanol without disturbing the pellet and the supernatant was again removed after 30 seconds. The ethanol washing step was repeated, and the tube was spun down and placed on the magnetic rack, with excess ethanol removed. The samples were air-dried, then resuspended in 26 µL of 1X TE Buffer (PanReac AppliChem, A8569,1000) by flicking and incubating at 50°C for 1 minute. The sample was then incubated at room temperature for an additional 5 minutes before repelling it on the magnet. The eluate was then removed and transferred into a 0.2 mL PCR tube (Sarstedt, 72.990.002) or 1.5 mL DNA LoBind tube (Eppendorf, 0030108051).

### Coriell Samples

Human genomic DNA samples were obtained from the Coriell Institute for Medical Research, as detailed in **Supplementary Table 14**. DNA concentration, purity, and DIN were measured as described above upon receipt.

### Library preparation and ONT Cas9-targeted sequencing

The library preparation was performed as in Erdmann et al.^8^ for R9 (R9.4.1) MinION flow cells, with some adaptations for R10 (R10.4.1) MinION flow cells. Upon arrival, guide RNAs (crRNA, IDT, **Supplementary Table 1**) and tracrRNAs (IDT, 1072533) were resuspended in TE-Buffer of pH 7.5 (IDT, 11-05-01-05) to 100 µM and stored at -80°C until further use. To produce the gRNA complex, 1 µL of each crRNA pool was mixed with 1 µL of tracrRNA in 8 µL of Duplex buffer (IDT, 11-01-03-01), annealed by heating at 95°C for 5 minutes, and cooled to room temperature. Next, 79.2 µL of nuclease-free water, 10 µL of rCutSmart™ Buffer (NEB, B6004), and 0.8 µL of Alt-R™ S.p. HiFi Cas9 Nuclease V3 (IDT, 1081060) were added to the annealed RNAs, forming the ribonucleoprotein (RNP) complexes, which were incubated at room temperature for 30 minutes. These RNPs were then placed on ice for further use. In parallel, 5 µg of input gDNA in 24 µL nuclease-free water were dephosphorylated by incubation with 3 µL Quick CIP Phosphatase (NEB, M0525) and 3 µL rCutSmart™ Buffer at 37°C for 20 min (adapted from the original protocol, which specifies a 10-minute incubation), followed by enzyme deactivation at 80°C for 2 min and kept at RT until further use. To the dephosphorylated DNA, 10 µL of the RNP complex, 1 µL of dATP (NEB, N0440), and 1 µL Taq DNA Polymerase (NEB, M0273) were added. The reaction mixture was incubated at 37°C for 15 min and 5 min at 72°C. The mix was then cooled to 4°C and stored on ice until further use. Adapter ligation was performed by combining 20 µL of ligation buffer (ONT, SQK-LSK114 kit), 3 µL of nuclease-free water, 10 µL NEBNext Quick T4 DNA ligase (NEB, E6056), and 5 µL adapter mix (ONT, SQK-LSK114 kit). Immediately after mixing, the adapter ligation mix was added to the cleaved and dA-tailed sample to yield an 80 µL final reaction volume. The resulting mixture was incubated at room temperature for 30 min (adapted from the original protocol, which specifies a 10-minute incubation), and 1 volume 1X TE pH 8.0 (IDT,11-01-02-05 11-01-02-5 was added to stop the reaction. DNA was purified using 48 µL AMPure XP beads (AXP) (ONT, SQK-LSK114 kit) and Long Fragment Buffer (LFB) (ONT, SQK-LSK114 kit) according to the manufacturer’s protocol, and the sample eluted in 13 µL Elution Buffer (EB) (ONT, SQK-LSK114 kit). To prime the flow cells, 1 ml Flow Cell Flush (FCF) (ONT, SQK-LSK114 kit) was combined with 25 µL Flow Cell Tether (FCT) (ONT, SQK-LSK114 kit), and 800 µl of the mix was slowly inserted in the Priming Port after removing any air bubbles. After a 5-minute incubation, the SpotON Sample Port was opened, and another 200 µl of the mix was added via the Priming Port. 12 µL of the library was mixed with 37.5 µL Sequencing Buffer (SB) (ONT, SQK-LSK114 kit) and 25.5 µL Library Beads (LIB) (ONT, SQK-LSK114 kit) before loading in a dropwise fashion onto the SpotON sample port of the primed flow cell (ONT, FLO-MIN114) for sequencing on the GridION Mk1 sequencer. The run was started after incubating the library on the flow cell for 10 minutes and placing the light shield on the sample port and sensor array. Ligation Sequencing Kit LSK114 and the Cas9 Sequencing Expansion were chosen to start the run, and the run limit was set to 30 hours.

### Multiplexing tests

#### Exonuclease and proteinase K treatments

The gRNA complexes, RNPs preparation, dephosphorylation, and Cas9 cleavage steps were performed as above with an input mass of 4µg as recommended by ONT to be sequenced on R10 (R10.3 flowcells). When no dephosphorylation was performed in the assay, 3 µL of rCutSmart Buffer (NEB, B6004S) and 3 µL nuclease-free water were added before the Cas9 digestion. For exonuclease treatment with Lambda Exonuclease (NEB, M0262S), 5 µL of Lambda Exonuclease Reaction Buffer (NEB, B0262S) was mixed with 1 µL of nuclease-free water. For the combination of Lambda Exonuclease and Exonuclease I (NEB, M0263S), 5 µL of Exonuclease I Reaction Buffer (NEB, B0293S) was mixed with 1 µL 1:200 diluted UltraPure BSA solution (Thermo Fisher Scientific, AM2616). To the samples, 4 µL of Lambda Exonuclease and 4 µL of Lambda Exonuclease with 1 µL of Exonuclease I were added after the addition of 6 µL custom buffer mix. After flicking the tube and centrifugation, the reaction was heated for 30 minutes at 37°C, then heated to 80°C for 5 minutes to deactivate the exonucleases. Samples treated with Thermolabile Proteinase K (NEB, P8111S) had one 1 µL of the protein added, were flicked, spun down, and heated to 37°C for 15 minutes and 80°C for 4 minutes to deactivate the enzyme. After the reaction, The volume was adjusted to 50 µL with 9 µL of nuclease-free water. Samples not treated with exonucleases and Proteinase K were heat treated at 80°C for 5 minutes to denature Cas9, then brought to a volume of 50 µL by adding 10 µL nuclease-free water and flicked. The dA-tailing was performed as in the library preparation above, with 1.5 µL of dATP (NEB, N0440) and 1.5 µL Taq DNA Polymerase (NEB, M0273) to match the increased volume but with the same temperature profile. 52.5 µL of AXP (ONT, SQK-NBD114.96) were added at a 1X ratio, and the samples were cleaned two times with 250 µl 80% EtOH without resuspending and eluted in 7.5 µL nuclease-free water.

### Multiplexing samples with the Ataxia panel

Samples were prepared as above with 4 µg DNA input, the protocol performed as before without exonuclease digestion, Proteinase K digestion, and dephosphorylation of the samples, and the dA-tailing performed as described before in the library preparation protocol. Additionally, the volume was not adjusted, and the clean-up was subsequently done with 42 µL of AMPure XP (Beckman Coulter, A63882) beads, 80% of EtOH, and eluted in 7.5 µL of nuclease-free water into a 0.2 mL PCR tube (Sarstedt, 72.990.002). Afterward, the protocol was continued with the ligation of barcodes.

### Ligation of barcodes for multiplexing tests

7.5 µL of the samples were mixed with 2.5 µL unique Native Barcode (NB01-96) from the Native Barcoding kit (ONT, SQK-NBD114.96) and 10µL Blunt/TA Ligase Master Mix (NEB, M0367), the sample mix flicked and spun down. After incubation for 20 minutes at room temperature, 2.0 µL of EDTA from the Native Barcoding kit (ONT, SQK-NBD114.96) was added, and the solution was mixed by flicking. Samples to be run together were then pooled in a 1.5 mL DNA LoBind tube (Eppendorf, 0030108051). Then AXP (ONT, SQK-NBD114.96) were mixed on the vortex, 9 µL per sample in the pool of the beads were added at a 0.4X ratio and incubated on a Thermomixer (Eppendorf, 5382000015) for 10 minutes at 450 rpm at room temperature. Samples were pelleted on a DynaMag™-2 Magnet magnetic rack (Thermo Fisher Scientific, 12321D) until the solution was clear. The supernatant was discarded, 250 µL Short fragment Buffer (SFB) (ONT, SQK-NBD114.96) was added, and the pellet was resuspended and spun down. The washing step was repeated, and the samples spun down and again placed on the magnetic rack to remove excess SFB, and allowed to air-dry for 30 seconds, then removed from the magnet, resuspended in 35 µL nuclease-free water, and were incubated for 10 minutes at 37°C on the Thermomixer (Eppendorf, 5382000015) at 450 rpm. The samples were again pelleted for at least 1 minute on the magnetic rack until the solution was clear and colorless, and 34 µL of the eluate was transferred to a 1.5 mL DNA LoBind tube (Eppendorf, 0030108051). Samples were optionally stored at this point at 4°C overnight.

### Ligation of adapters, priming, and loading for multiplexing tests

To the now pooled samples, 10 μL 5X NEBNext Quick Ligation Buffer, 5 μL NEB T4 DNA Ligase from the NEB Quick Ligation Module (NEB, E6056), and 1.0 μL Native Adapter (NA) ONT, SQK-NBD114.24) were added, the tubes were flicked and spun down, then incubated for 20 minutes at room temperature. AXP (ONT, SQK-NBD114.96) were vortexed, 20 μL per sample was added at a 0.4X ratio and incubated on a Thermomixer (Eppendorf, 5382000015) for 10 minutes at 450 rpm at room temperature. Samples were pelleted on a DynaMag™-2 Magnet magnetic rack (Thermo Fisher Scientific, 12321D) until the solution was clear. The supernatant was discarded, and 250 μL LFB (ONT, SQK-NBD114.24)/SFB (ONT, SQK-NBD114.24) was added, the samples were removed from the magnet, the pellet was resuspended and spun down. The washing step was repeated, samples spun down, and excess LFB was removed; the samples were allowed to air-dry for 30 seconds, then removed from the magnet, resuspended in 15 μL Elution Buffer (ONT, SQK-NBD114.24), and incubated for 10 minutes at 37°C on the Thermomixer (Eppendorf, 5382000015) at 450 rpm. The samples were again pelleted for at least 1 minute on the magnetic rack until the solution was clear and colorless, and 12 μL eluate was transferred to a 0.5 mL DNA LoBind tube (Eppendorf, 0030108035).

This library was mixed with 37.5 μL SB and 25.5 μL (LIB) in the tube and stored on ice until applied in the flow cell. Then, 1170 μL of the FCF was mixed with 30 μL Flow Cell Tether (FCT) (ONT, SQK-NBD114.24) and 5 μL UltraPure BSA solution (Thermo Fisher, AM2616). After ensuring no air bubbles were present in the priming port of the flow cell by extracting a small volume, 800 μL of the above mix was gently loaded into the priming port of the R10 GridION flow cell (ONT, FLO-MIN114), incubated for 5 minutes and then the remaining 200 μL of priming buffer was loaded into the flow cell after the sample spot port had been opened. Afterward, 75 μL of the sample was added dropwise into the sample port, and the sequencing run started after 5 minutes. The Native Barcoding Kit 96 was selected, a sequencing duration of 12 to 72 hours was selected depending on the experiment, and the run started after placing the light shield on the sample port. The sequencing time was 30 hours, 48 hours, or 72 hours for the ATX panel and 12 hours for the multiplexing tests.

### Reloading flow cells

To reuse flow cells after sequencing, the Flow Cell Wash Kit (ONT, EXP-WSH004) was used as per the manufacturer’s instructions. Briefly, 2 µL of the Wash Mix (WMX) was mixed with 398 µL of the Diluent (DIL) (ONT, EXP-WSH004) and placed on ice. For rerunning the library, the sequencing run was paused, and the old library was removed by extracting 150 µL volume from the priming port and storing it in a 0.5 mL DNA LoBind tube (Eppendorf, 0030108035). After ensuring no air bubbles were present in the sample port, 200 µL of the prepared mix was added to the flow cell through the priming port and incubated for 5 minutes. The rest of the mix was loaded into the flow cell and incubated for 1 hour. The flow cell was then either filled with 500 µL Storage Buffer (S) (ONT, EXP-WSH004) and stored at 4°C for further use or primed again using the flow cell priming mix, and 150 µL of the extracted library was reloaded into the sample port.

### Agarose gels

1,000-4,000 ng of DNA was combined with a running buffer of 40% sucrose in 100 mL of distilled water with Orange G. The 1% agarose gels were prepared by combining 1% w/v Agarose Type II-A (Sigma Aldrich, A9918-250G) in 1X TBE consisting of 16.35% Trizma-base (Sigma Aldrich, 93352), 2.78% boric acid (Sigma Aldrich, B0252-500g), 0.93% EDTA (Sigma Aldrich, E5134-500G) and 2X MidoriGreen advanced (NIPPON Genetics, MG04). The gel was prepared in a Sub-Cell GT Cell (Bio-Rad, 1704401), and the DNA samples were run in 1X TBE buffer with HyperLadder 2 kb (meridian, BIO-33053) at 80V until separated (approx. 90-120 minutes). The gels were imaged using a UV gel imaging system (Nippon and Tuppon GLS) with an exposure time between 0.3 and 2 seconds.

### Analysis of Oxford Nanopore sequencing data

Basecalling and methylation from POD5 files were performed using Dorado (v0.6.2), and the resulting FASTQ files mapped to the human using Minimap2 (epi2me-labs/wf-alignment v1.1.0)^23^ to identify the reads spanning the targets of interest. NanoPlot was used For quality control of the aligned reads (v.1.29.1)^24^.

#### Multiplexing runs

Sequencing data was processed using Epi2ME v5.1.14 and the Epi2ME agent v3.7.3 provided by ONT. The output included the proportion of barcodes in the sequencing run and the .csv file containing data on the aligned reads. For further analysis, specifically to create BAM files that were used for the coverage analysis in IGV, the Epi2ME desktop version was used to align reads to the hg38 genome. The workflow used was either the wf-alignment, which aligns reads to the chosen genome or the wf-Cas9 pipeline, which outputs coverage information based on a target gene BED file. The output files were sorted into barcodes, and the coverage was read out by loading them into IGV and parsing them into a .csv file that could then be further processed with the statistical tool R (4.4.0). The pore count data was taken from the run reports, which also provided information on the data output of the flow cell.

## DATA AVAILABILITY

Anonymized basecalled POD5 nanopore data will be deposited to the European National Archive (ENA) upon publication. From this data, both basecalled POD5 and/or FASTQ files can be acquired.

## ACKNOWLEDGEMENTS

We thank the patients who provided samples for this study and for providing informed written consent, as well as Isabel Abellan Schneyder from ONT for support and input on the project.

## AUTHOR CONTRIBUTIONS

V.Schönrock performed the majority of the lab work for R10 validation. V.Scholz performed the bioinformatic analysis of the R10 validation data. VP performed the lab work and analysis for the Cas9 multiplexing tests. HE, AA, EHF, and TN provided clinical expertise and devised the gene panels. M.Schoedel and MA validated the guide RNAs. HE, MCL VM, EB, IvB, and CD assessed the sequencing data and clinical results. MCL, ABP, AA, and EHF conceived the project. MCL and ABP supervised the work. EHF, AA, and TN provided the funding and resources to conduct the work. MCL, V.Scholz, and VP prepared the figures. MCL wrote the paper, with contributions from all authors.

